# Deciphering the evolution of microbial interactions: in silico studies of two-member microbial communities

**DOI:** 10.1101/2022.01.14.476316

**Authors:** Gayathri Sambamoorthy, Karthik Raman

**Affiliations:** Robert Bosch Centre for Data Science and Artificial Intelligence (RBCDSAI), Indian Institute of Technology (IIT) Madras, Chennai – 600 036, INDIA; Centre for Integrative Biology and Systems mEdicine (IBSE), IIT Madras, Chennai – 600 036, INDIA; Department of Biotechnology, Bhupat and Jyoti Mehta School of Biosciences, IIT Madras, Chennai – 600 036, INDIA

## Abstract

Microbes thrive in communities, embedded in a complex web of interactions. These interactions, particularly metabolic interactions, play a crucial role in maintaining the community structure and function. As the organisms thrive and evolve, a variety of evolutionary processes alter the interactions among the organisms in the community, although the community function remains intact. In this work, we simulate the evolution of two-member microbial communities *in silico* to study how evolutionary forces can shape the interactions between organisms. We employ genome-scale metabolic models of organisms from the human gut, which exhibit a range of interaction patterns, from mutualism to parasitism. We observe that the evolution of microbial interactions varies depending upon the starting interaction and also on the metabolic capabilities of the organisms in the community. We find that evolutionary constraints play a significant role in shaping the dependencies of organisms in the community. Evolution of microbial communities yields fitness benefits in only a small fraction of the communities, and is also dependent on the interaction type of the wild-type communities. The metabolites cross-fed in the wild-type communities appear in only less than 50% of the evolved communities. A wide range of new metabolites are cross-fed as the communities evolve. Further, the dynamics of microbial interactions are not specific to the interaction of the wild-type community but vary depending on the organisms present in the community. Our approach of evolving microbial communities *in silico* provides an exciting glimpse of the dynamics of microbial interactions and offers several avenues for future investigations.

## 1 Introduction

Microbes co-exist in complex communities, forming networks by virtue of their interactions with one another [1, 2]. These interactions are crucial for the existence of microbes in communities and also for performing essential functions of the ecosystem [3]. The nature and extent of interactions among organisms in the community determine community function [4]. Therefore, the emergent properties that arise out of community relationships cannot be understood by studying microbes in isolation but rather by considering the interactions among the microbes. The interactions can be positive, negative, or neutral. Interactions among microbes are typically classified into six types, viz. amensalism, commensalism, mutualism, neutralism, parasitism, and competition [1].

The co-evolution of organisms in microbial communities plays a central role in the existence and survival of microbes in a community [5, 6]. Microbes in a community coordinate with each other, conserving community functions [7]. Studies show that the structure and diversity of microbial communities are influenced largely by evolutionary processes [8–10]. Microbes respond to various selection pressures, assimilating genetically driven changes over time. As a result, the associations among the organisms in the community are altered, although the community may remain intact. It is interesting to study how the community functions are preserved while organisms acquire new capabilities via evolutionary processes. Understanding the evolution of interactions that exist among microbes in a community is an open question in microbial ecology [11, 12].

Selective forces play a significant role in maintaining the community structure and altering the interaction patterns among the organisms in a community [13]. Further, individual species selection is inadequate for engineering microbial communities for a particular application, such as the sustained production of a product of interest or enhancing stress-tolerant community interactions. Often, a selection at the community level is desired rather at the individual level while studying organisms in a community. For example, artificial selection experiments have been performed based on above-ground plant biomass [14]. Experiments have been performed to identify the artificial selection methods to improve the functions of the community [15]. Further, it is interesting to study how different selection pressures favour certain interactions.

In this work, we seek an evolutionary understanding of microbial communities from a metabolic perspective since microbes primarily interact by exchanging extracellular metabolites. We employ a Markov Chain Monte Carlo (MCMC) approach to evolve two-member microbial communities comprising organisms from the human gut, which exhibit varied interaction patterns. Next, we study how the microbial interactions evolve as the organisms in the community have a change in the genetic architecture. We examined the growth phenotypes of the microbial communities upon evolution to study if evolution confers fitness benefits. We also analysed the metabolites cross-fed among organisms as they evolve. Finally, we study the dynamics of microbial communities comprising arbitrary microbes, as compared to naturally-existing communities of the human gut. Overall, our study paves the way for understanding the evolution of metabolic interactions in microbial communities and their role in shaping community structure.

## 2 Methods

### 2.1 Choice of organisms for evolution of microbial communities

We chose organisms from the human gut to simulate the evolution of naturally existing microbial communities. We selected 20 organisms from the Virtual Metabolic Human (VMH) database (https://vmh.life/; [16]), which spanned the four major phyla in the human gut, namely, Firmicutes, Bacteroidetes, Proteobacteria, and Actinobacteria. Supplementary File S1 contains the list of these 20 organisms.

We simulated all possible pairwise community models from these 20 organisms. For each of these 20C_2_ communities, we identified those that could grow together, on a ‘ Western diet’ [17], and predicted their interaction types. The communities were first grouped based on their interaction types, and from each interaction type, we chose five example communities for further study. The list of communities selected and their growth rates in the starting community are given in the Supplementary File S1.

Further, we picked a few additional organisms for studying co-evolution, from the BiGG knowledgebase (https://bigg.ucsd.edu/; [18]). These organisms were chosen such that the organisms, when simulated together in a community, could sustain growth. The list of the BiGG models employed for analysis and the communities formed from these eight organisms, and the interaction types they exhibit are tabulated in Supplementary File S2.

### 2.2 Evaluation of growth phenotypes in microbial communities

The joint models (‘community metabolic network’) for the two-member microbial communities were created using the `createMultiSpeciesModel` routine from the COBRA toolbox [19]. Further, the growth phenotypes were predicted using community FBA (cFBA) [20]. cFBA is a constraint-based approach for modelling microbial communities and predicting growth phenotypes. cFBA employs linear programming for maximising the sum of the biomass fluxes of the organisms in the community. The formulation of cFBA is as given below:

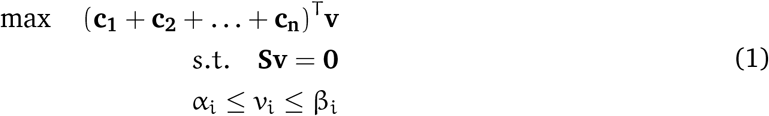

where **c_1_**, **c_2_**, …, **c_n_** are objective functions that correspond to the biomass reaction of *n* individual organisms, **v** is the vector of fluxes of all the biochemical reactions in all organisms, including the exchange reactions of all the cross-feeding metabolites, and **S** corresponds to the stoichiometric matrix of the reactions present in the combined network of all *n* organisms, again including the exchange reactions. α_i_ and β_i_ signify the lower and upper bounds of the flux of the i^th^ reaction in the community metabolic network.

### 2.3 Constraints for community evolution

To study how the community dynamics change for varied evolutionary pressures, we employed three different selection constraints. Microbial communities perform a specific function of the ecosystem and evolve to retain the functionality. Here, we consider the overall growth of the community as the function of the microbial community. We devised three different constraints based on the overall community growth, which is the sum of each organism’s growth when present in a community. The first constraint, 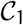, demands no selection based on community growth. In contrast, the second constraint 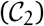 requires minimal community growth, while the third constraint 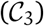 enforces a higher community growth. A minimum fraction of their individual growth (the growth of each organism when present individually) is a common requirement in all three cases. The mathematical formulation of the constraints is depicted in Table 1.

**Table 1:** Constraints employed for community evolution. The table lists the three constraints employed for community evolution and the growth that is demanded in each constraint. v_1,com_ and v_2,com_ are the growth rates of Organism 1 and 2 when present in the community. v_1,ind_ and v_2,ind_ are the growth rates of Organism 1 and 2 respectively, when simulated individually. 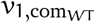, 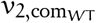, 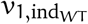, and 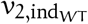 are the fluxes of the Organisms 1 and 2 in the community and when present individually in the wild-type, *i.e.* the microbial community at the start.

| List of constraints | Constraints employed |
| --- | --- |
| **Constraint 1 ($\mathcal{C}_1$)** | $v_{1,\text{ind}} > 0.1 v_{1,\text{ind}_{WT}}$ |
| | $v_{2,\text{ind}} > 0.1 v_{2,\text{ind}_{WT}}$ |
| **Constraint 2 ($\mathcal{C}_2$)** | $v_{1,\text{com}} > 0.1 v_{1,\text{com}_{WT}}$ |
| | $v_{2,\text{com}} > 0.1 v_{2,\text{com}_{WT}}$ |
| | $v_{1,\text{ind}} > 0.1 v_{1,\text{ind}_{WT}}$ |
| | $v_{2,\text{ind}} > 0.1 v_{2,\text{ind}_{WT}}$ |
| **Constraint 3 ($\mathcal{C}_3$)** | $v_{1,\text{com}} > 0.5 v_{1,\text{com}_{WT}}$ |
| | $v_{2,\text{com}} > 0.5 v_{2,\text{com}_{WT}}$ |
| | $v_{1,\text{ind}} > 0.1 v_{1,\text{ind}_{WT}}$ |
| | $v_{2,\text{ind}} > 0.1 v_{2,\text{ind}_{WT}}$ |

### 2.4 Evolving microbial communities ***in silico***

Individual organisms have been evolved *in silico*, by using an MCMC approach previously [21, 22]. In this approach, organisms are ‘evolved’ in every step, by means of reaction swaps. That is, beginning with the metabolic network of a given organism, at each iteration, one reaction in the organism is swapped out with another reaction from a ‘universe’ of possible reactions. If the organism remains viable (*i.e.* shows growth on the given medium) post-swap, the swap is retained, completing one iteration of the evolution. Otherwise, the ‘failed’ swap is discarded and the network is reverted to its previous state. This approach has been used to study the evolvability of metabolic networks [21] and redundancy [22].

Here, we extended this approach to model the evolution of two-member microbial communities. We took the ‘universe’ set of reactions as a union of all the reactions present in the VMH database. We pooled reactions from every organism in the database and removed the exchange reactions, biomass reactions, sink reactions, and also reactions pertaining to replication, transcription, and protein biosynthesis. For every microbial community under study, we used an iterative reaction swap approach (as illustrated in Figure 1), where one reaction is selected at random from either of the organisms in the community, and replaced with a randomly chosen reaction from the universe of reactions described above. Following this reaction swap, the resultant community is simulated to evaluate the constraints (Table 1). Once the constraints are met, the process is repeated iteratively. If the constraint is not met at any step, the resultant community is discarded, and the iteration is restarted with the community generated in the previous step. One evolved microbial community is generated out of 1, 000 such successful reaction swaps. We believe this approach captures key facets of the evolutionary forces that operate in nature, as has also been described previously [21].

**Figure 1:**
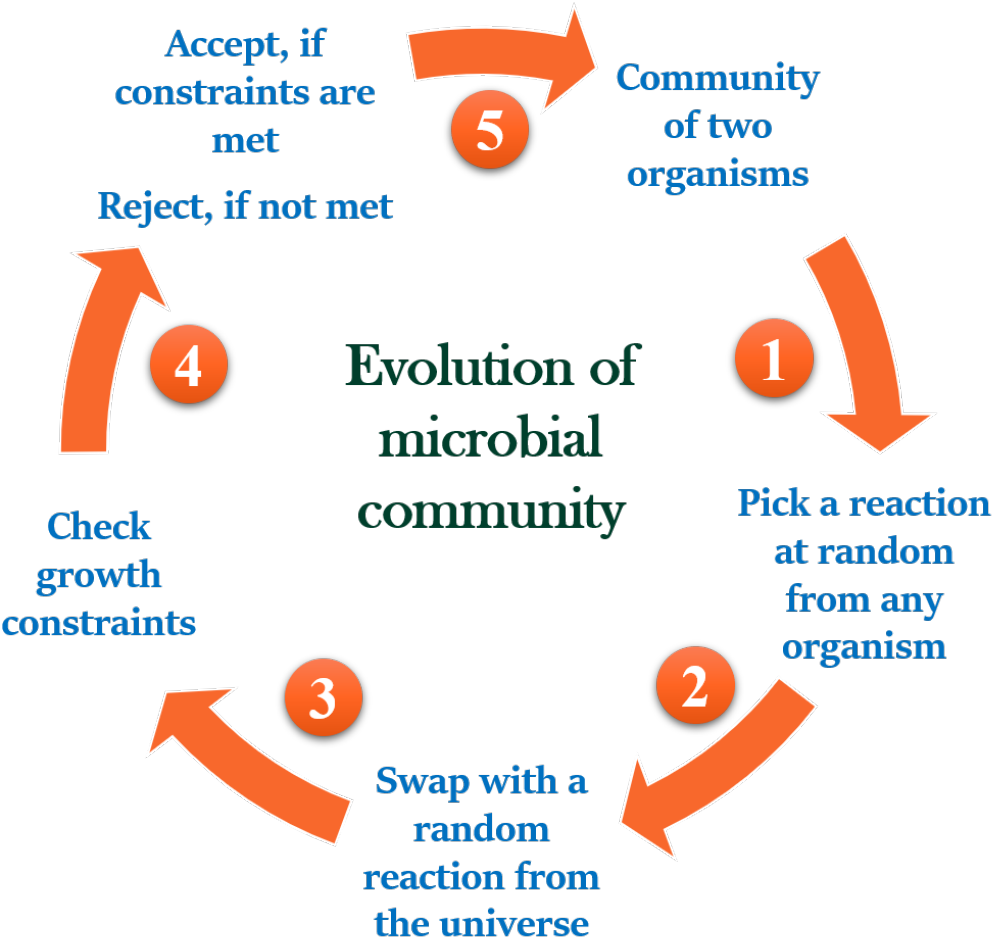
Schematic of the MCMC approach to evolve microbial communities *in silico*. The figure outlines the steps employed for evolving microbial communities using an MCMC approach. Beginning with a community of two organisms, one reaction is picked at random from either organism, swapped with a reaction randomly sampled from the universe of possible reactions, followed by checking if the constraints are satisfied to either accept the swap or reject (and repeat the process). This entire process is iterated 1, 000 times to produce one evolved community, for a given starting pair of organisms.

### 2.5 Identification of interaction type in communities

The interactions in microbial communities can be classified into six types: amensalism, commensalism, parasitism, mutualism, competition, and neutralism [1]. For identifying the interaction type in a given community, we adapt a previously defined approach of comparing growth rates of the organisms in the community with the growth rates when the organisms are allowed to grow individually [17]. Based on the growth rates of the organisms when present individually and when present as a community, we introduce two parameters α_1_ and α_2_ based on the growth rates, as:

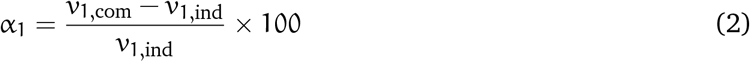

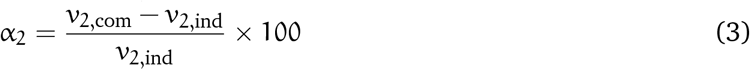

where v_1,com_ is the growth rate of Organism 1 in the community, v_1,ind_ the growth rate of Organism 1 when present alone. Similarly, v_2,com_ and v_2,ind_ are the growth rates of Organism 2 when present in a community and individually, respectively.

A cut-off of 10% is set to distinguish the communities into various interaction types. For example, α_1_ value of more than 10 indicates that Organism 1 is benefited from the community relationship. Similarly, an α_1_ value of less than −10 demonstrates that Organism 1 has a substantially reduced growth when present in the community. Based on the values of α_1_ and α_2_, the interaction types are classified as shown in Table 2.

**Table 2:** Classification of communities into different interaction types. The table depicts the classification of microbial communities into different interaction types based on the values α_1_ and α_2_. The shorthand notation readily indicates the nature of interaction—a zero means a non-significant (*i.e.* < 10% change), while a ‘+’/’–’ indicates a significant (*i.e.* > 10%) increase or decrease in the growth rates (α values), respectively.

| Interaction type | Shorthand | Range of $\alpha_1$ | Range of $\alpha_2$ |
| --- | --- | --- | --- |
| Amensalism | $\{0, -\}$ | $-10 \leq \alpha_1 \leq 10$ | $\alpha_2 \leq -10$ |
| | $\{-, 0\}$ | $\alpha_1 \leq -10$ | $-10 \leq \alpha_2 \leq 10$ |
| Commensalism | $\{0, +\}$ | $-10 \leq \alpha_1 \leq 10$ | $\alpha_2 \geq 10$ |
| | $\{+, 0\}$ | $\alpha_1 \geq 10$ | $-10 \leq \alpha_2 \leq 10$ |
| Competition | $\{-, -\}$ | $\alpha_1 \leq -10$ | $\alpha_2 \leq -10$ |
| Mutualism | $\{+, +\}$ | $\alpha_1 \geq 10$ | $\alpha_2 \geq 10$ |
| Neutralism | $\{0, 0\}$ | $-10 \leq \alpha_1 \leq 10$ | $-10 \leq \alpha_2 \leq 10$ |
| Parasitism | $\{-, +\}$ | $\alpha_1 \leq -10$ | $\alpha_2 \geq 10$ |
| | $\{+, -\}$ | $\alpha_1 \geq 10$ | $\alpha_2 \leq -10$ |

The organisms in the community are further classified based on the community relationship. An amensal community comprise two organisms—*amensal affected* and *amensal unaffected*; while amensal affected loses fitness benefits, amensal unaffected does not have any (substantial) fitness change when present in a community. Commensals comprise *commensal taker*, the organism in the community that gains fitness benefits, and *commensal unaffected*, the organism in the community that does not have any change in growth. Parasitic communities comprise a *parasitism giver*, the organism in the community that has a reduction in growth, and *parasitic taker*, the organism that acquires growth benefits.

### 2.6 Jaccard similarity

Jaccard similarity measures how similar two sets are [23]. We employed Jaccard similarity as a measure of the similarity of two organisms, based on their constitutive reactions. The Jaccard similarity is calculated as 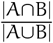 , where |A ∩ B| denotes the number of reactions common between the two organisms A and B, and |A ∪ B| denotes the total number of unique reactions present in both organisms.

### 2.7 Identification of cross-fed metabolites

The metabolites that are cross-fed between a pair of organisms are identified by analysing the flux distributions generated as a result of cFBA. The flux of the exchange reactions for each organism in the community is compared. If one organism has a positive flux (which signifies the secretion of the metabolite) and the other organism has a negative flux (which signifies the uptake of the metabolite) for a particular metabolite, then that metabolite is considered as cross-fed. While comparing the fluxes, a minimum cut-off flux of 0.01 mmol gDCW^−1^ h^−1^ was used for the identification of cross-fed metabolites.

## 3 Results

Organisms endure changes in their genetic architecture upon adaptations. For organisms that exist in microbial communities, how do such changes, modelled in terms of the addition/removal of reactions affect the nature of interactions and the existence of the community? Do interactions among the organisms in communities remain intact upon evolution? Does evolution alter the interaction type among the organisms? How do the metabolic dependencies among organisms in a community vary as they evolve? To address such questions, we examined the evolution of several microbial communities *in silico*.

### 3.1 Stability of microbial communities upon evolution

To evaluate the stability of a community, *i.e.* the sustained existence of the species in the community, we chose 20 organisms in the human gut from the VMH database and formed two-member communities (see Methods §2.1) that exhibited different interaction types (see Methods §2.5). The evolution was simulated *in silico* using an MCMC approach as described in §2.4. 1, 000 such evolved communities were generated for the 30 different example two-member communities under study (a total of 30 × 1000 evolved communities). The communities were evolved with a selection pressure that each organism that occur in a two-member community should sustain minimal growth when simulated individually (using 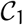 as mentioned in §2.3). Further, the evolved communities were simulated using cFBA and the steady-state fluxes were predicted. The biomass (*i.e.* growth rate) of each organism in the evolved communities was analysed.

Interestingly, we found that there are cases where the organisms thrive individually but do not sustain growth in the community. For each of the 1, 000 evolved communities for every example studied here, we identified the number of communities where either of the organisms showed zero growth when simulated using cFBA. For each interaction type, we identified the fraction of evolved communities in which at least one organism has non-zero growth. The fraction of 1000 evolved networks was averaged across the five example communities for each interaction type (Figure 2). The figure illustrates that the commensal wild-type communities yield more *stable* communities upon evolution than other interaction types. One would expect mutualistic communities to be stable owing to the ability of organisms to mutually benefit each other. However, we found that even parasitic communities (that manifest a predator–prey relationship), on average, had a higher number of communities that had one organism showing non-zero growth rate upon evolution. Unsurprisingly, competitive communities showed a fraction of 0.74 out of the evolved communities with either one organism with non-zero growth and the lowest among all interaction types, followed by amensal communities.

**Figure 2:**
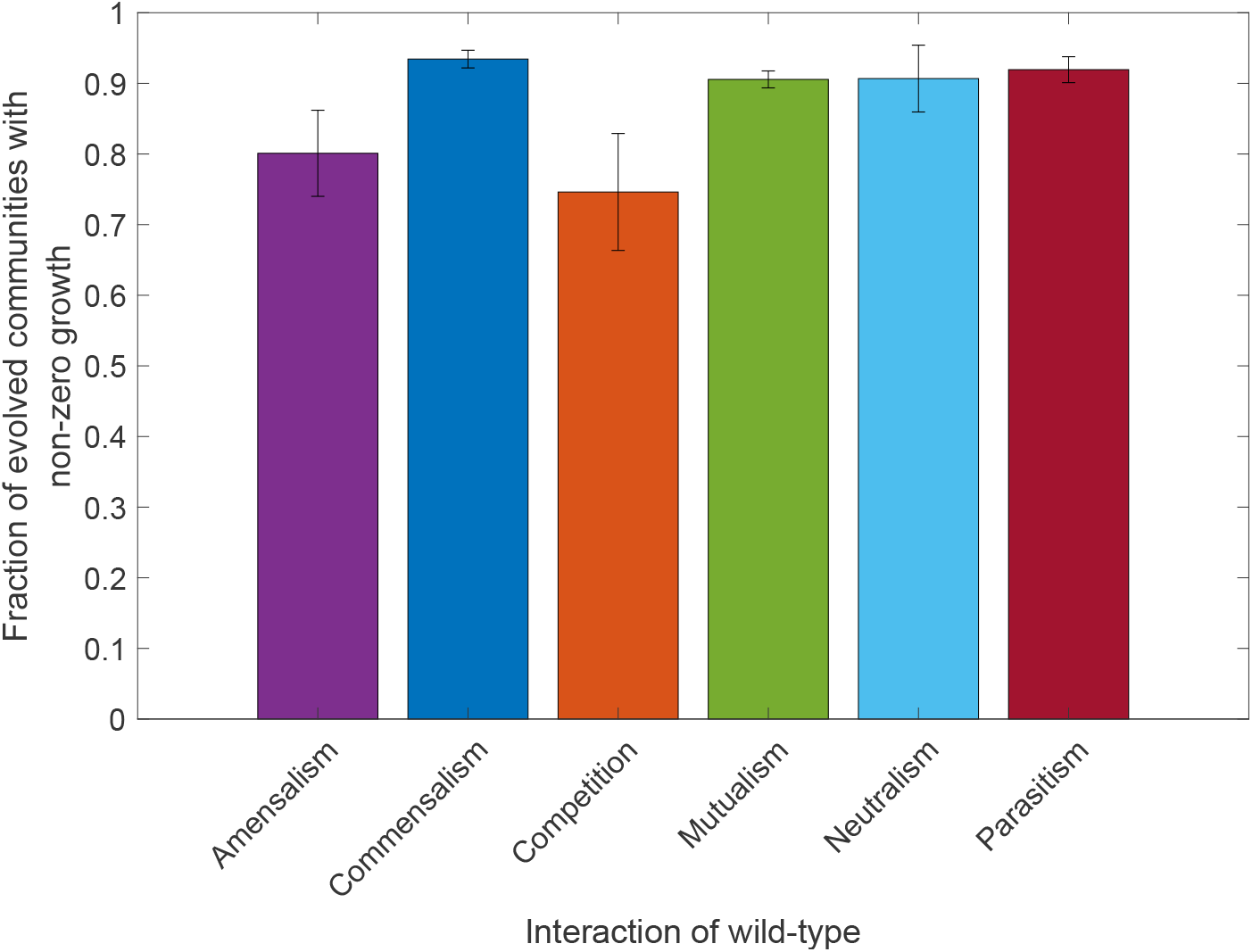
Growth in evolved communities. The bar describes the number of two-member evolved communities out of 1000 evolved communities in which either of the organism has zero growth. The number is averaged for all the examples pertaining to each interaction type of the wild-type community. The error bars indicate one standard deviation.

Further, we also studied the fraction of evolved networks in which an organism yields non-zero growth upon evolution and the same is tabulated in Table S1 in Supplementary Results for different interaction types. Table S1 shows that in some cases, a single organism in the community predominates, thereby, has a lower number of cases where there is non-zero growth in the community. Most examples for mutualistic and neutralistic communities had one organism that produces non-zero growth in the community upon evolution. On a closer look, we found that the *Ec-Ec*^1^ pair (in wild-type competitive community) showed almost a competing number of cases where either organism pertains to non-zero growth. While analysing the organisms present in the microbial community, we found that the two organisms have higher similarity in terms of the reactions present in the network than the other examples (measured by calculating the Jaccard similarity of the two organisms in the microbial community as given in §2.6). This signifies that organisms that have similar metabolic capabilities tend to compete. The Jaccard similarity values calculated for all the examples can be found in Supplementary File S1.

In the case of commensal and parasitic communities, one would expect that the organism that acts as the ‘giver’ thrives to a greater extent than the ‘taker’ in the community upon evolution. However, upon evolution, in commensal communities, the giver does not thrive in all cases. Although, in parasitic communities, in all cases, we find that the giver predominates the other organism present in the two-member community.

In summary, evolution alters the co-existence of organisms in two-member communities, and this particularly varies depending on the interaction type exhibited by the organisms. Organisms that are more similar in terms of the metabolic capabilities tend to compete with each other in the community, while one organism predominates the other in the case of organisms that have different reactions.

### 3.2 Evolution leads to varied interaction types

Next, we sought to understand how interactions transform as microbes evolve in communities. Upon evolution of a community belonging to a particular interaction type, what kind of interactions do the evolved communities exhibit? Are interaction patterns among organisms that exist in communities *persistent*? To answer these questions, we employed the metabolic networks evolved using constraint 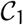 from each example community, which spanned across different interaction patterns. By simulating the communities using FBA and cFBA, we determined the growth rates of the organisms individually and in the community. Using the parameter values α_1_ and α_2_ calculated based on the growth rates, we identified the interaction type of each of the 1, 000 evolved communities for every example under study (see §2.5). Supplementary File S3 contains the fraction of evolved networks that exhibit different interaction patterns for every example studied across all the wild-type interaction categories. All amensal and neutralistic wild-type communities yield a major fraction of parasitic communities, followed by commensals, upon evolution. Commensal, mutualistic, competitive, and parasitic communities, although, upon evolution, produce parasitic communities in majority of the cases. In all evolved networks, the fraction of competitive communities appeared to be the lowest across the different interaction patterns of the wild-type communities.

We further investigated how the switch in the interaction behaviour is influenced by the constraints imposed during evolution. For the same, we set up experiments for evolution by varying the constraints for evolution. We employed two other constraints, 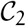 and 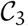, which additionally impose a threshold on the combined growth of the organisms in the community, besides individual growth. For each of the constraints, we evolved 1, 000 networks for every example community, similar to the previous experiment. For all the evolved networks, we identified the interaction patterns and found the average interaction behaviour of the communities evolved from every wild-type interaction pattern (average of the five examples for each wild-type interaction pattern). Figure 3 depicts the fraction of networks that exhibit different interaction patterns in the evolved networks and the changes in the interaction patterns that are affected due to the change in the constraint.

**Figure 3:**
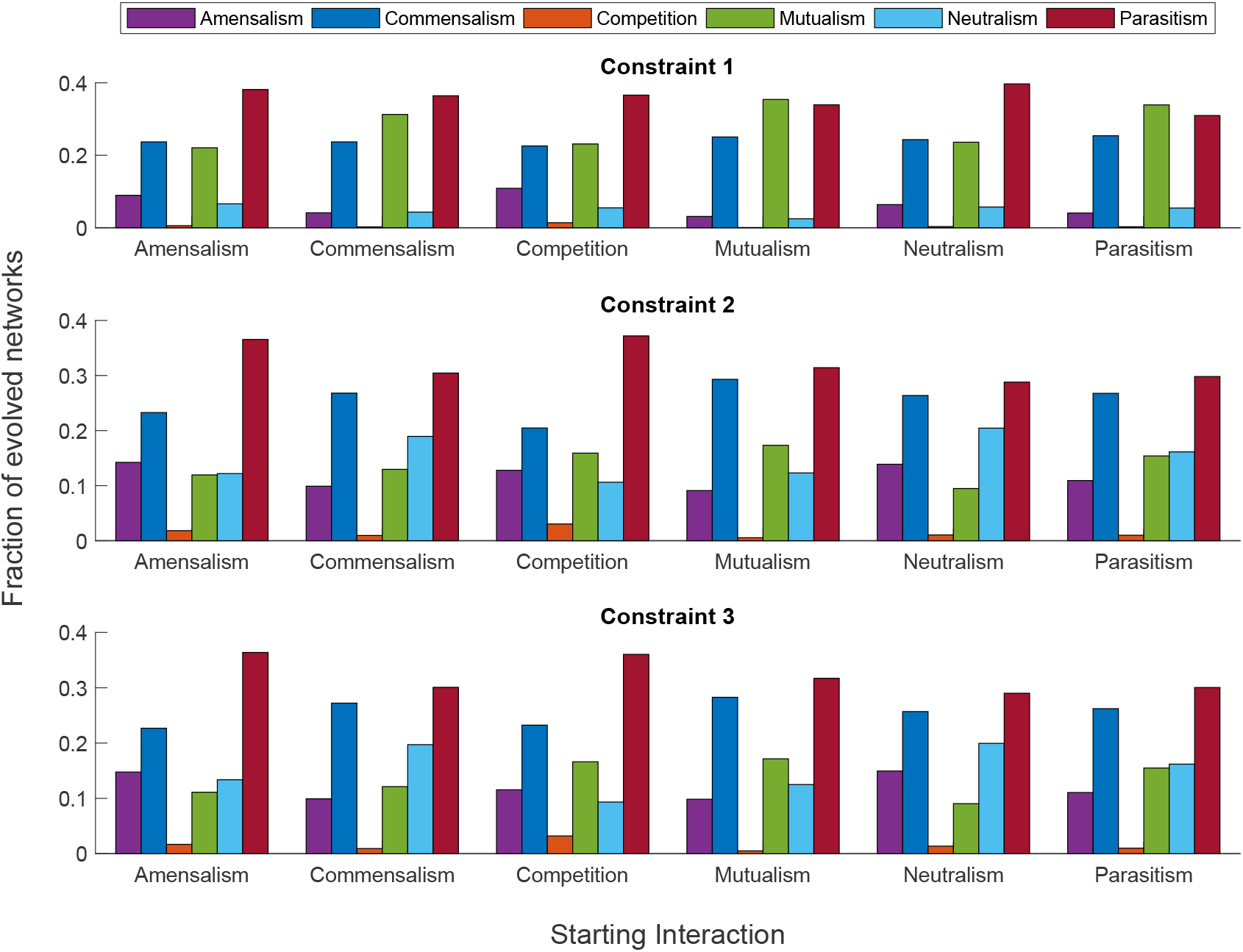
Change in interaction patterns upon evolution on different constraints. The figure shows the fraction of evolved communities that exhibit different interaction patterns. The evolved communities are grouped based on the interaction type of the wild-type communities.

On analysis of the communities evolved using constraints 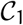 and 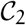, we observed a similar trend in the interaction types exhibited by organisms. Among all the interactions, parasitic communities were predominant in the evolved communities, irrespective of the interaction type of the wild-type community. Following parasitism, we observed that commensals form the next prominent among the evolved communities.

The constraint 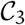 yields a major fraction of parasitic interaction in the communities evolved from amensal, commensal, competition, and neutralistic wild-type communities. Interestingly, on the other hand, mutualistic and parasitic communities, upon evolution, yield a larger fraction of mutualistic behaviour. Competitive behaviour was rarely observed in the communities that were evolved using constraint 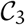, with mutualistic wild-type communities showing almost zero competitive communities, upon evolution.

To conclude, evolution leads to communities exhibiting varied interactions. Importantly, this suggests that the community interactions are not *stable* — a mixture of interaction types is observed in the evolved communities. The constraints play a significant role in altering the interaction patterns of the organisms in the communities. An important finding is that the fraction of communities that evolved into mutualistic behaviour were higher in networks evolved using constraint 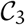 than those observed in constraints 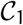 and 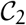.

In essence, evolution drives organisms in communities to alter the interactions that exist among them. Interactions among organisms in a two-member community is not persistent and vary depending on the reactions constituted in the organisms.

### 3.3 Fitness benefits depends on the nature of interactions

It is interesting to understand how the growth patterns of the community change upon evolution. Does evolution increase the growth rate in microbial communities? Does the starting interaction have an impact on how the community acquires growth benefits? Does the growth of the evolved community depend on what type of interactions the organisms exhibit? One may expect that upon evolution, since organisms are forced into a community relationship, the growth rate of the community, v_com_ would increase compared to their mono-cultures. We computed the ratio of predicted community growth to their predicted growth in mono-cultures for all evolved communities. We compared the ratio with the growth ratios in the wild-type. The difference in the ratios between the evolved communities and the wild-type communities was found and plotted in Figure 4. The figure describes the change in community growth upon evolution grouped by the interaction pattern of the wild-type communities. All evolved communities have an average decrease in community growth.

**Figure 4:**
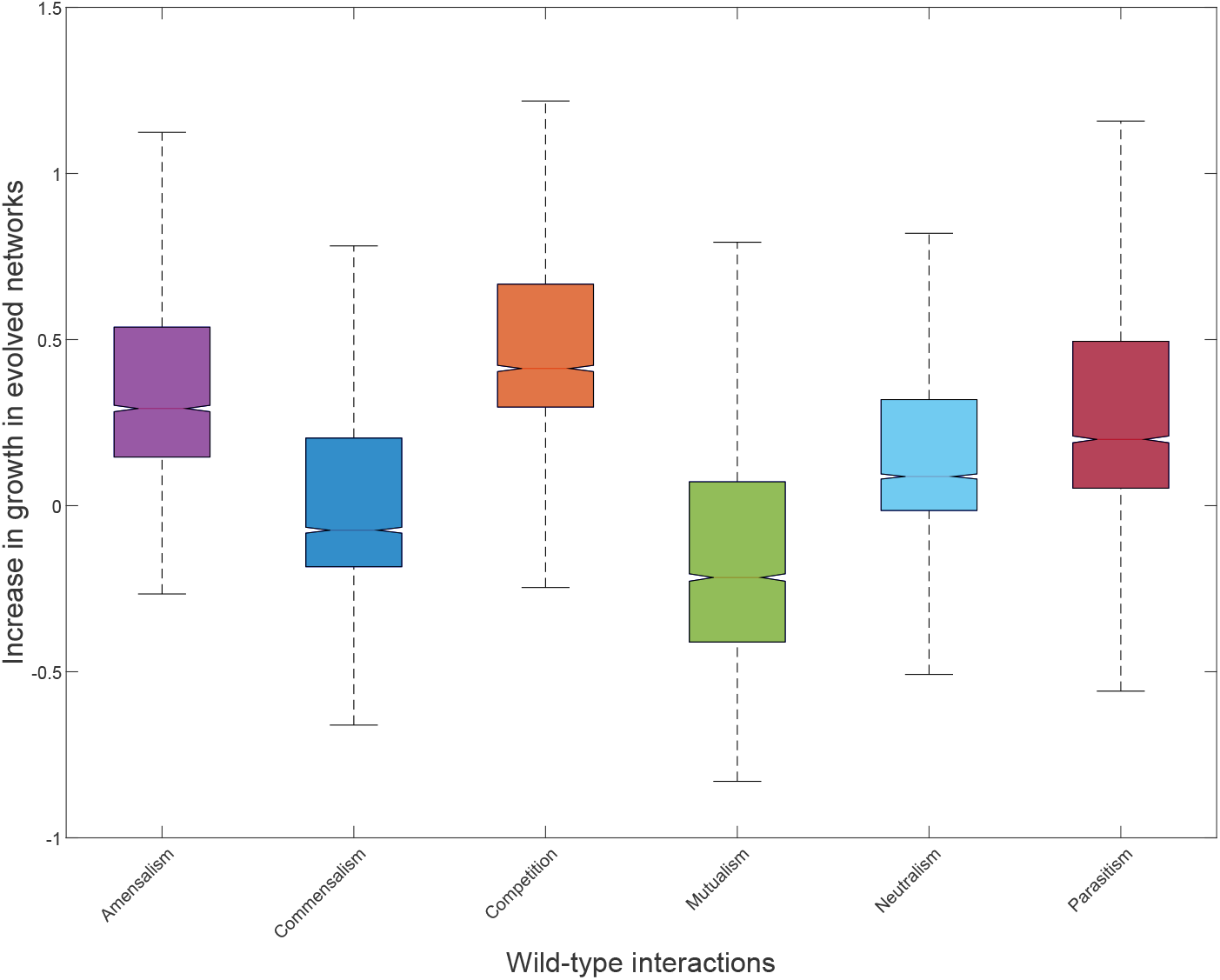
Change in community growth upon evolution. The figure depicts the change in community growth upon evolution. The distribution shows the difference in ratios grouped based on the interaction type of the wild-type communities.

The mean of the distribution was higher for competitive communities with a value of 0.532. This indicates that the communities evolved from competitive wild-type communities showed higher growth in the community than the mono-cultures. Interestingly, communities evolved from mutualistic wildtype communities showed the lowest mean of −0.0758. Followed by competitive wild-type communities, evolution from amensal communities yielded a distribution with a higher mean increase in the ratio of growth rates in co-cultures and mono-cultures.

Does evolution confer fitness benefits, measured in terms of the overall growth of the community? For the analysis, we identified the percentage of evolved communities that exhibit a higher growth than the wild-type communities. Particularly, we were interested in identifying the percentage of communities that show high growth as they evolve from one interaction to the other. We grouped the percentage of high-growth communities according to the wild-type interaction and the evolved interaction. The heat map in Figure 5 depicts the percentage of evolved networks that exhibit higher growth compared to the wild-type, as they evolve from one interaction type to the other. Communities that evolved from wild-type that exhibited amensal behaviour tend to show a larger percentage of networks that have high growth than other wild-type communities. On the other hand, commensal wild-type communities displayed the least percentage of evolved networks exhibiting high growth. Around 10% of the communities that were evolved from amensal to parasitism had higher growth than the wild-type. Interestingly, the communities that were evolved from mutualistic wild-type communities showed the least percentage of networks that manifest higher community growth.

**Figure 5:**
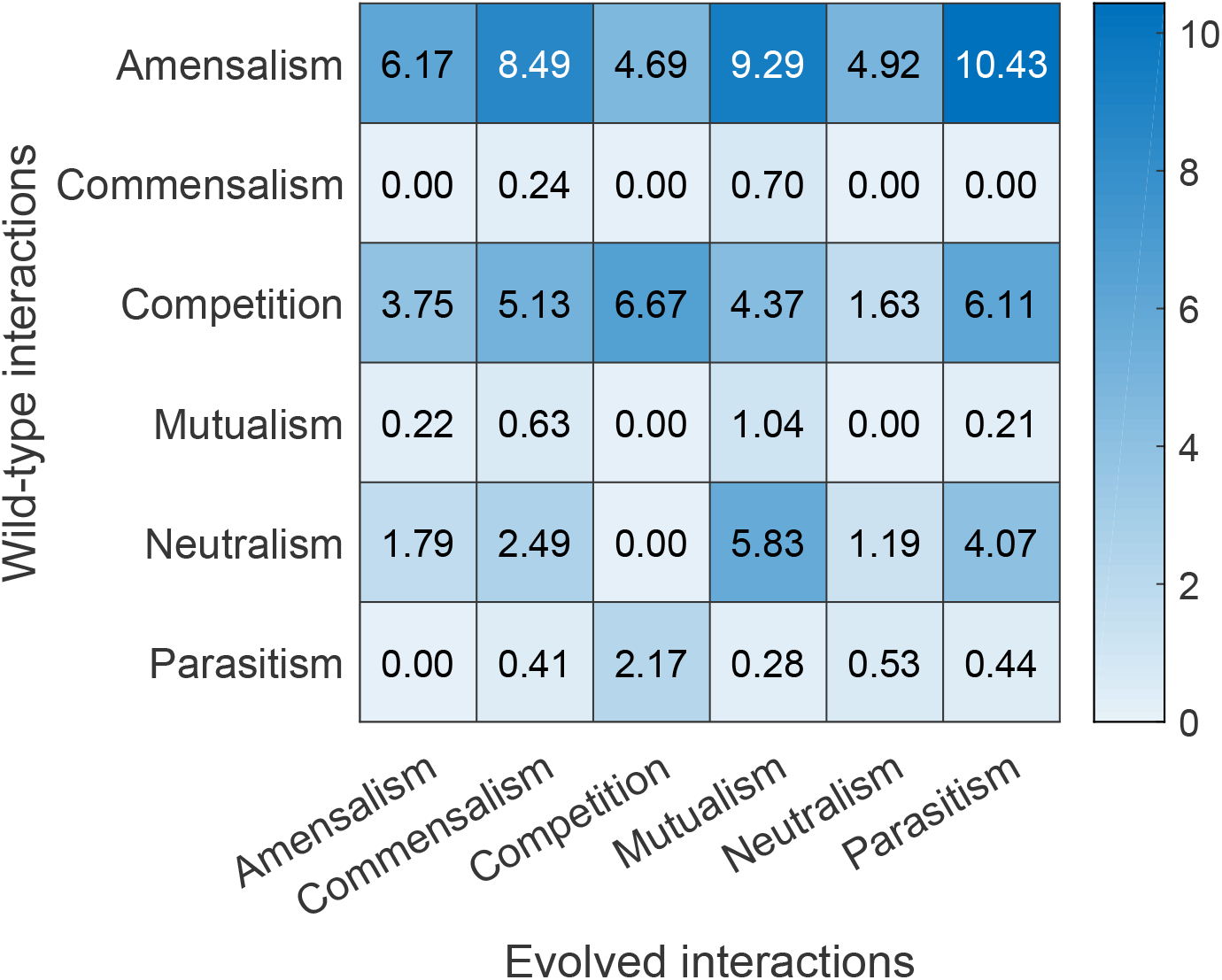
Percentage of high-growth evolved communities. The heat map shows the percentage of high-growth evolved communities that exists when they evolve from one interaction type to the other. The rows pertain to the wild-type interaction while the columns denote the evolved interaction type.

In sum, the growth benefits from a community relationship upon evolution vary with the interaction type of the starting communities. A small fraction of the evolved communities exhibited fitness benefits compared to the wild-type communities. The growth benefits of the organisms in the evolved communities vary depending on the interactions exhibited by the wild-type community and also on the interaction type they evolve into.

### 3.4 Evolution results in varied metabolic dependencies

Next, we studied the metabolic exchanges among organisms in the two-member microbial communities under study. These metabolites are termed as cross-fed, *i.e.* they are secreted from one organism and utilised by the other. Initially, we identified the cross-fed metabolites in the wild-type communities using the flux distributions obtained when simulated using cFBA (as described in §2.5). Similarly, we found the cross-fed metabolites for all the communities evolved using 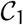. We compared the number of metabolites cross-fed in the initial community and the evolved communities. Figure 6 shows the distribution of the number of metabolites cross-fed in the evolved communities for all the examples under study and the number of metabolites cross-fed in the wild-type communities. We found that the number of metabolites exchanged in the wild-type was always the same or higher than the mean of the number of metabolites exchanged in evolved communities.

**Figure 6:**
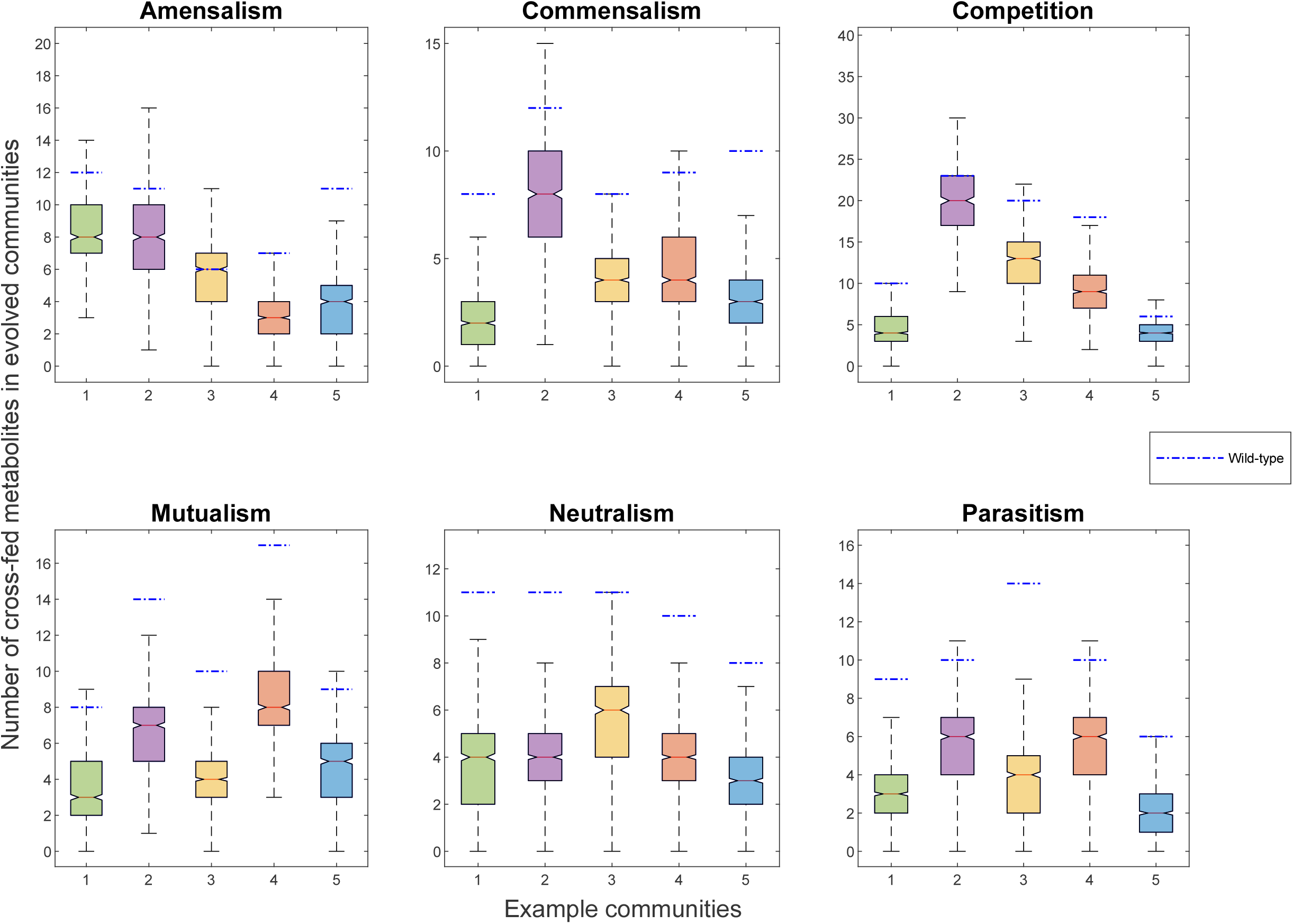
Number of metabolites cross-fed in evolved communities. The figure shows the distribution of the number of cross-fed metabolites in evolved communities. The blue dashed line indicates the number of exchanges in the wild-type community.

Additionally, we investigated if the metabolic exchanges that exist in the wild-type are retained when the communities evolve. The wild-type communities exchange varied metabolites, and the number of times these exchanges were observed in the evolved networks was found. In all cases, the cross-fed metabolites were observed in only less than 50% of the evolved networks. The number of times each cross-fed metabolite is observed across the 1000 evolved communities for each example is documented in Supplementary File S4.

Further, we analysed the cross-fed metabolites in the evolved communities. We observed that amino acids were predominant in the metabolites cross-fed in the evolved communities, such as leucine, lysine, glutamine, aspartate, and also, in some cases other metabolites such as succinate, glycolaldehyde, and ornithine. The metabolite that is maximally exchanged in each example community for all interaction types is shown in Table S2 in Supplementary Results.

Thus, the metabolic dependencies in communities vary as the communities evolve. In the majority of the cases, amino acids are cross-fed in the evolved communities. On the whole, the cross-fed metabolites depend on the nature of the organisms in the community and the metabolic dependency that emerge as a result of evolution.

### 3.5 Synthetic communities behave markedly different from natural communities

We next study if synthetic communities behave differently from communities comprising organisms that co-exist in nature. For this, we chose arbitrary organisms that are not known to occur in a community naturally and assembled them together to analyse their community behaviour. For this, we selected eight models from BiGG (as listed in §2.1) and predicted the growth of their pairwise community models. We further short-listed those communities in which both organisms demonstrate growth. For all these communities, we predicted the interaction types exhibited by the organisms. The list of 21 communities used in the study is tabulated in Supplementary File S2. All the communities were further subjected to evolution, with minimal growth of 10% of the wild-type growth. 1, 000 evolved communities were generated for each of the 21 communities, as described in Methods §2.4.

We predicted the interaction patterns of the evolved communities. The fraction of each interaction type exhibited was plotted for every interaction pattern of the wild-type communities, as shown in Figure 7. We observed that a major fraction of co-evolved communities showed amensal behaviour, while communities evolved from parasitic and mutualistic wild-type communities exhibited a significant fraction of commensals. It is interesting to note that amensals and commensals are interactions where one organism in the community is neither benefited nor harmed. Further, our study suggests that communities evolving into mutualistic relationships are minimal in most of the wild-type interaction types.

**Figure 7:**
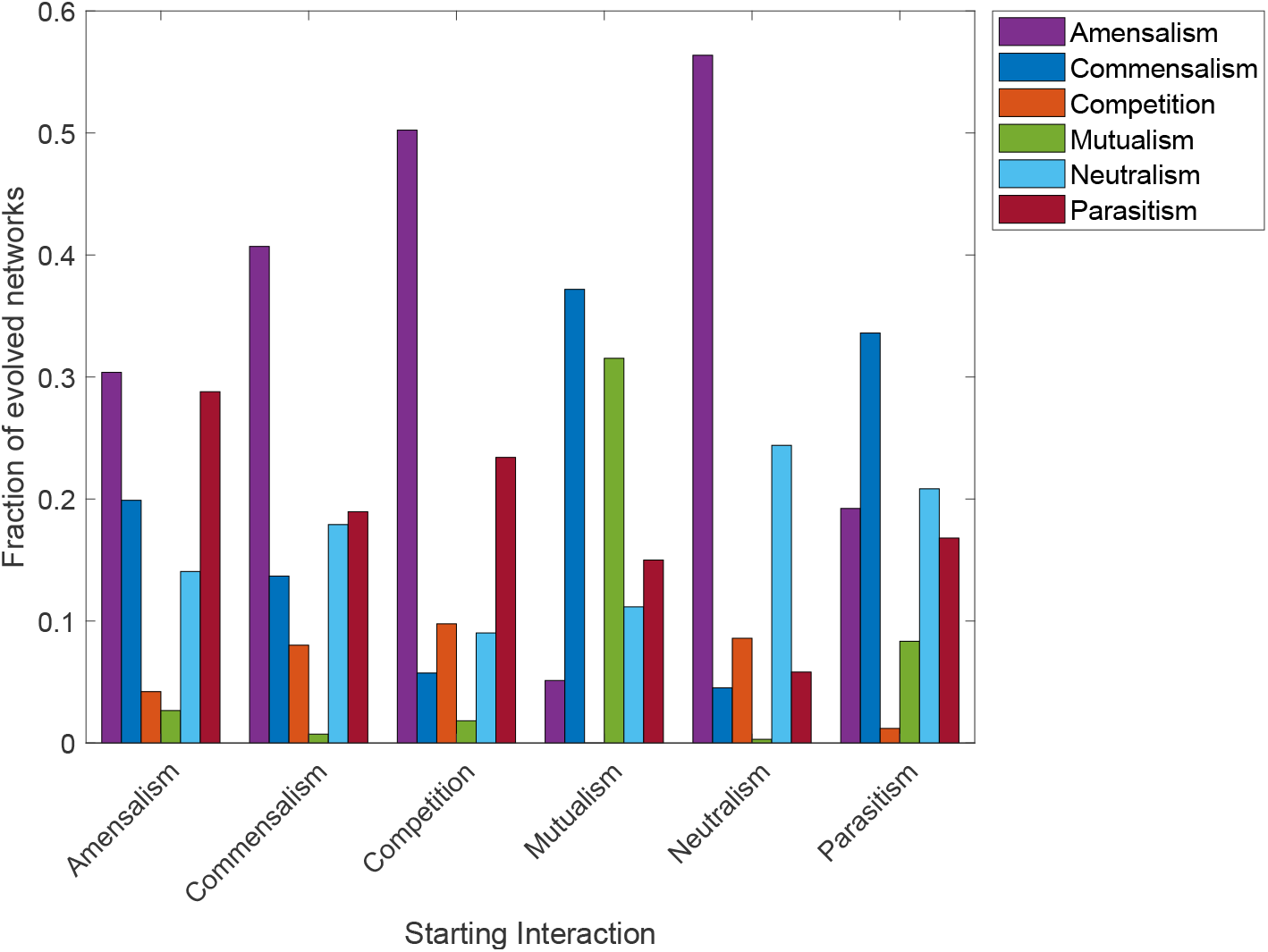
Change in interaction patterns upon evolution of arbitrary communities. The figure shows the fraction of evolved communities that exhibit different interaction patterns. The evolved communities are grouped based on the interaction type of the wild-type communities.

Thus, we observe that the interaction patterns in the evolved communities for the arbitrary communities are different from those found in the communities of the human gut. It is evident that the dynamics of the microbial communities vary for natural communities as opposed to ‘arbitrary’ microbial communities.

### 3.6 Interactions in the evolved communities depends on the organisms in the starting community

We earlier studied (§3.2) the interaction patterns of the evolved communities in order to comprehend the changes in interaction as organisms evolve. The interaction type exhibited by organisms in a community is predicted by calculating the α_1_ and α_2_ values for all the evolved communities. Since a threshold of 10% is employed while identifying the interaction patterns, we chose to analyse the α values across the evolved communities. Does altering the cut-off have an influence on the classification of the interaction types? We also aimed to study the above under the three different constraints employed. We identified the α_1_ and α_2_ values of the different co-evolved communities as described in §2.5. The α_1_ and α_2_ values were discretised into different bins and were plotted as a heat map along with the wild-type α values as depicted in Figure 8. The figure depicts the plot of α values for all evolved networks from each wild-type community exhibiting amensalism and for different constraints. The figure illustrates the distribution of evolutionary outcomes, showing the ‘extent’ of the relationship evolved. We found that different starting communities exhibit varied interaction patterns in the evolved communities. Examples *Kp-Bo* and *Bt-Cb* evolved using 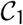 show more parasitic behaviour than the other examples that show a majority of mutualistic behavior.

**Figure 8:**
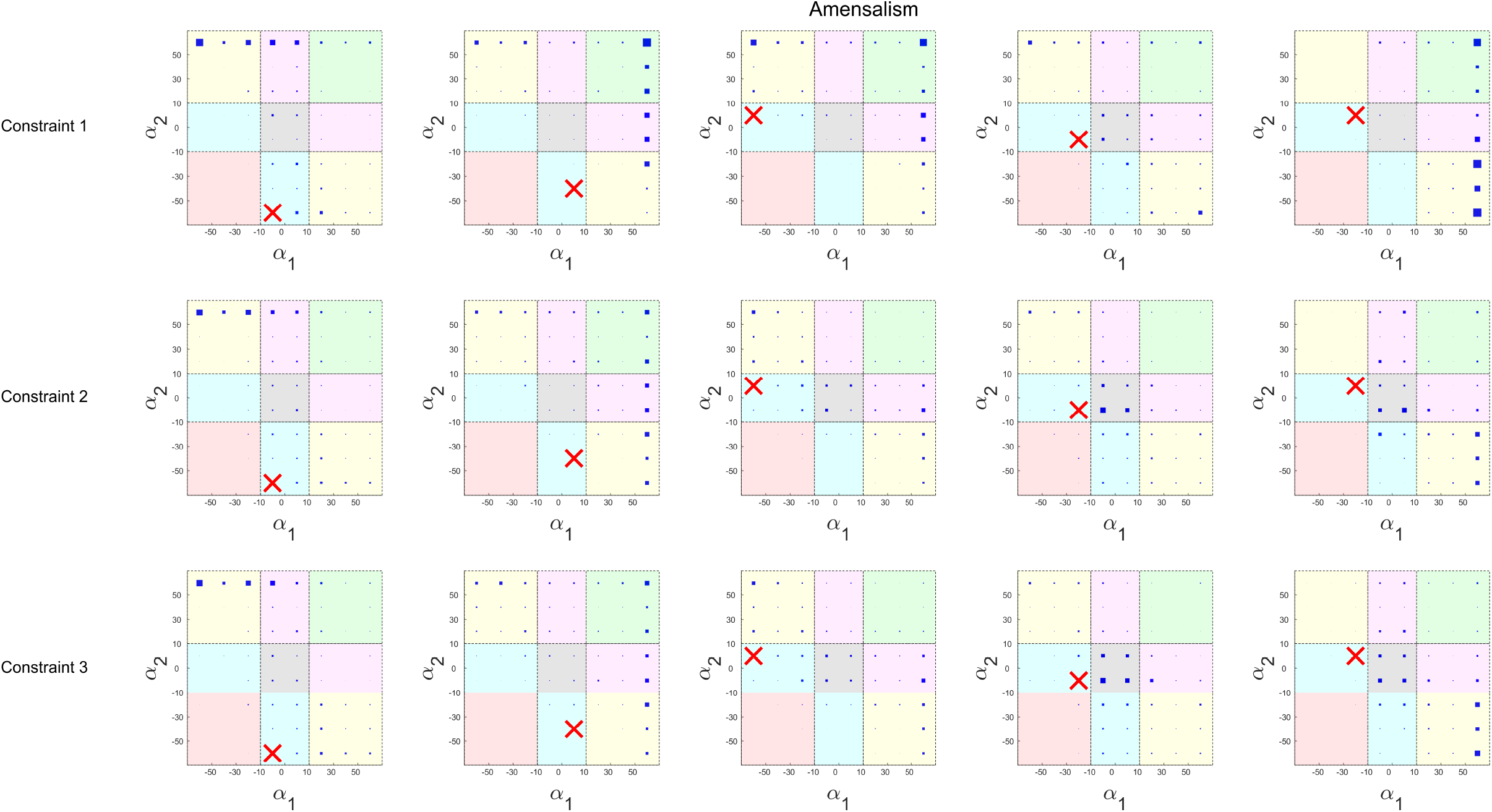
Heat map of α_1_ and α_2_ values for amensal starting communities. The figure depicts the density of α_1_ and α_2_ in each bin for all the evolved networks from a starting community that exhibits amensal behaviour. Each subplot denotes the evolved communities from an example starting community. The different rows represent the constraints. The graph is coloured into regions based on the cut-off of 10% into various interaction types as green-mutualism, pink-commensalism, yellow-parasitism, blue-amensalism, grey-neutralism, orange-competition. The red cross denotes wild-type starting community.

Further, while comparing across constraints, we find that constraints 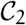 and 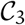 show similar outcomes, as opposed to 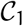. When we compare how the α values have evolved from the starting communities, we find that the α values do not have a significant role to play in how the networks evolve. This is because we observe that the bin of the starting community does not determine the fate of the α values in the evolved communities. An interesting observation is that the values of α_1_ and α_2_ of the evolved communities are highly concentrated on the edges of the histogram, indicating that the threshold does not alter the outcome of the interaction types drastically. This also shows that the communities exhibit a higher propensity to evolve into a particular interaction type that is mostly dependent on the nature of the organisms in the community. A similar plot was illustrated for all the starting communities exhibiting different interaction patterns. They are depicted in Figures S1–S5 in the Supplementary Results.

In sum, we studied the extent of α values for each of the evolved communities for every wild-type community. We found that the individual communities vary in how the interaction patterns evolve. Constraints show a change in the α values, although constraints 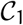 and 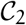 yielded similar results. Thus, the way in which communities evolve depends on the organisms present in the community and is not unique to communities exhibiting a particular interaction pattern.

## 4 Discussion

Microbes co-operate and co-exist in communities by the exchange of metabolites. For biotechnological applications, the study of microbial communities is on the rise [24–29], with several studies focussing on the evolutionary stability and performance of communities [12, 30–35]. Understanding the evolutionary dynamics of the interactions exhibited by the communities is thus interesting. Here, we employ metabolic models for studying the *in silico* evolution of communities that show varied interaction patterns. Such metabolic modelling analysis can help understand the generic nature of evolutionary mechanisms, which is not possible or time-consuming by employing experiments.

First, we explored the stability of organisms in two-member microbial communities. Communities are not always *stable*, in that the organisms upon evolution do not necessarily continue to exist in a community relationship. Previous evolution experiments conducted on independent co-cultures led to the extinction of species in some evolved communities [30]. Next, we analysed the evolutionary trajectory of communities, with different initial ‘configurations’, viz. mutualism, neutralism, amensalism, commensalism, parasitism or competition. Our *in silico* evolution experiments predict that the interactions between a given pair of organisms is necessarily not stable, and a variety of end-points are possible. In most evolved communities, the proportion of competitive interaction was lower than other interaction types. Previous studies also suggest that although organisms tend to compete for resources initially, the level of competition reduces locally, forming stable communities [36].

The fitness of the evolved communities was dependent on the interaction of the wild-type communities. While analysing the overall fitness, calculated as the sum of growth rates of both organisms in the community, a small fraction of the evolved communities showed an increase in community growth. Perhaps, a different selection constraint that demands the coordination of both organisms in the community would yield higher fitness benefits. Nevertheless, when microbes co-ordinate in natural communities to perform an essential function of the ecosystem, some organisms might get knocked out due to other organisms’ acquired abilities to meet the purpose. This observation is in accordance with a prior study, where the evolution of co-cultures in a uniform environment led to higher biomass in some cases, while some did not show any improvement [30].

Ideally, one would expect microbes in a community to evolve such that they are beneficial for one another, although that is not what we observe. Organisms in communities exhibit varied cross-feeding metabolites as communities evolve. A recent modelling study shows that evolutionary contingency determines the evolution of cross-feeding in microbial communities [37]. Similarly, we found that some communities exhibit a higher number of metabolite exchanges, while many evolved communities have lower cross-fed metabolites. Similar to a study that showed that metabolic dependencies arise although they are not selected for [38], we found a range of metabolic dependencies among organisms in the community. Evolution does not necessarily lead to cooperative networks, considering the cost of secretion of metabolites, leading to single species producing all the metabolites that it requires [33]. Our analysis showed that the majority of the metabolites exchanged among the evolved communities were amino acids that are known to occur for co-operation among microbial communities in some previous studies [33, 39].

We found that organisms present in the community play a significant role in the interaction patterns exhibited by the evolved communities. Further, the constraints play a major role in shaping the dependencies among organisms in the evolved communities. Our study also throws light on the differences in the way organisms present in natural communities evolve over arbitrary communities.

Our study does have some limitations. First, our study focuses on understanding the evolution of microbial interactions only by means of metabolic exchanges. However, other modes of interactions do exist that affect the evolution of dependencies among organisms in communities. However, metabolic exchanges have been shown to be very important drivers of community metabolism [2]. Further, the abundance of species also plays a crucial role in the dynamics of the interactions; these have not been accounted for in our modelling paradigms. Although pairwise interactions are crucial for the community structure, there exist higher-order interactions in microbial communities [40, 41]. Further, in a community composed of more than two species, complex principles that drive community structure exists, such as functional redundancy [42]. Therefore, unravelling the evolution of higher-order interactions will further our understanding of the emergent properties and dynamics in microbial communities. Nevertheless, pairwise interactions have indeed shown to be the major drivers of community dynamics, rather than higher-order interactions [43]. We have striven to capture selection pressures that exist in nature, by means of the three constraints 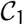, 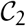, and 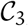. While it is difficult to capture the exact selection pressures/evolutionary forces at play in nature, these constraints provide a scenario analysis of possible outcomes in nature.

The approach employed here opens up several vistas for future investigations. For instance, our approach can be used to predict the stability of a microbial community in industrial fermentations. By defining additional constraints such as, say, the production of a metabolite of interest, our approach can be used to analyse if the communities could potentially evolve to produce more of a given product without compromising on the overall growth of the community. The dynamics of microbial interactions in varied nutrient conditions will further our understanding of the stability of microbial interactions. Overall, our unique approach provides an exciting glimpse of the evolutionary dynamics in microbial communities, paving the way for understanding the evolution of microbial interactions in communities, and other facets of community evolution, such as metabolic dependencies.

## Supporting information

Supplementary Results

## Supplementary Material

**Supplementary Results** contains the list of Supplementary tables and figures;

**Table S1** depicts the fraction of evolved networks with non-zero growth for organisms in the community for different interaction types.

**Table S2** lists the metabolites that are majorly exchanged in evolved communities for every example in each interaction type.

**Figures S1-S5** shows the spread of α1 and α2 values for networks evolved using the three different constraints for each starting interaction type.

**Supplementary File S1** contains the list of 20 organisms chosen from the VMH database. It also contains the example communities for each interaction type and the growth rates upon simulation.

**Supplementary File S2** lists the 21 models chosen from the BiGG database for simulation. It also lists the example communities for each interaction type and the growth rates upon simulation of the community.

**Supplementary File S3** denotes the fraction of evolved networks that exhibit each interaction type evolved using different constraints.

**Supplementary File S4** lists the metabolites cross-fed in the wild-type communities and frequency of their occurrence in the evolved networks.

## Footnotes

1 See Supplementary File S1 for communities and their two-letter codes

