## Supplementary Results for "Deciphering the evolution of microbial interactions: in silico studies of two-member microbial communities"


### Supplementary Material

Table S1: **Frequency of zero growth in community**. The table depicts the fraction of evolved networks each organism shows non-zero growth when simulated in a community. The table shows the data for all evolved communities for the five different examples studied, in each interaction type.

**Fraction of evolved networks with non-zero growth in either of the organism**

| **Interaction** | **Example 1** | | **Example 2** | | **Example 3** | | **Example 4** | | **Example 5** | |
|---|---|---|---|---|---|---|---|---|---|---|
| **Type** | **Org1** | **Org2** | **Org1** | **Org2** | **Org1** | **Org2** | **Org1** | **Org2** | **Org1** | **Org2** |
| **Amensalism** | 0.34 | 0.09 | 0.02 | 0.09 | 0.02 | 0.19 | 0.11 | 0.06 | 0.08 | 0.01 |
| **Commensalism** | 0.06 | 0.04 | 0.05 | 0.01 | 0 | 0.09 | 0.06 | 0 | 0.02 | 0.01 |
| **Competition** | 0.01 | 0.2 | 0.31 | 0.27 | 0.16 | 0.02 | 0.14 | 0 | 0.06 | 0.11 |
| **Mutualism** | 0.08 | 0 | 0.10 | 0.01 | 0 | 0.05 | 0.11 | 0.01 | 0 | 0.11 |
| **Neutralism** | 0.05 | 0 | 0.06 | 0 | 0.24 | 0.04 | 0.03 | 0 | 0 | 0.04 |
| **Parasitism** | 0.02 | 0.01 | 0.13 | 0.01 | 0.07 | 0 | 0.08 | 0 | 0.06 | 0.04 |

Figure S1: **Heat map of $\alpha_1$ and $\alpha_2$ values for commensalism starting communities.** The figure depicts the density of $\alpha_1$ and $\alpha_2$ in each bin for all the evolved networks from a starting community that exhibits commensal behaviour. Each sub-graph denotes the evolved communities from an example starting community. The different rows represent the constraints. The graph is coloured into regions based on the cut-off of 10% into various interaction types as green-mutualism, pink-commensalism, yellow-parasitism, blue-amensalism, grey-neutralism, orange-competition. The red cross denotes wild-type starting community.

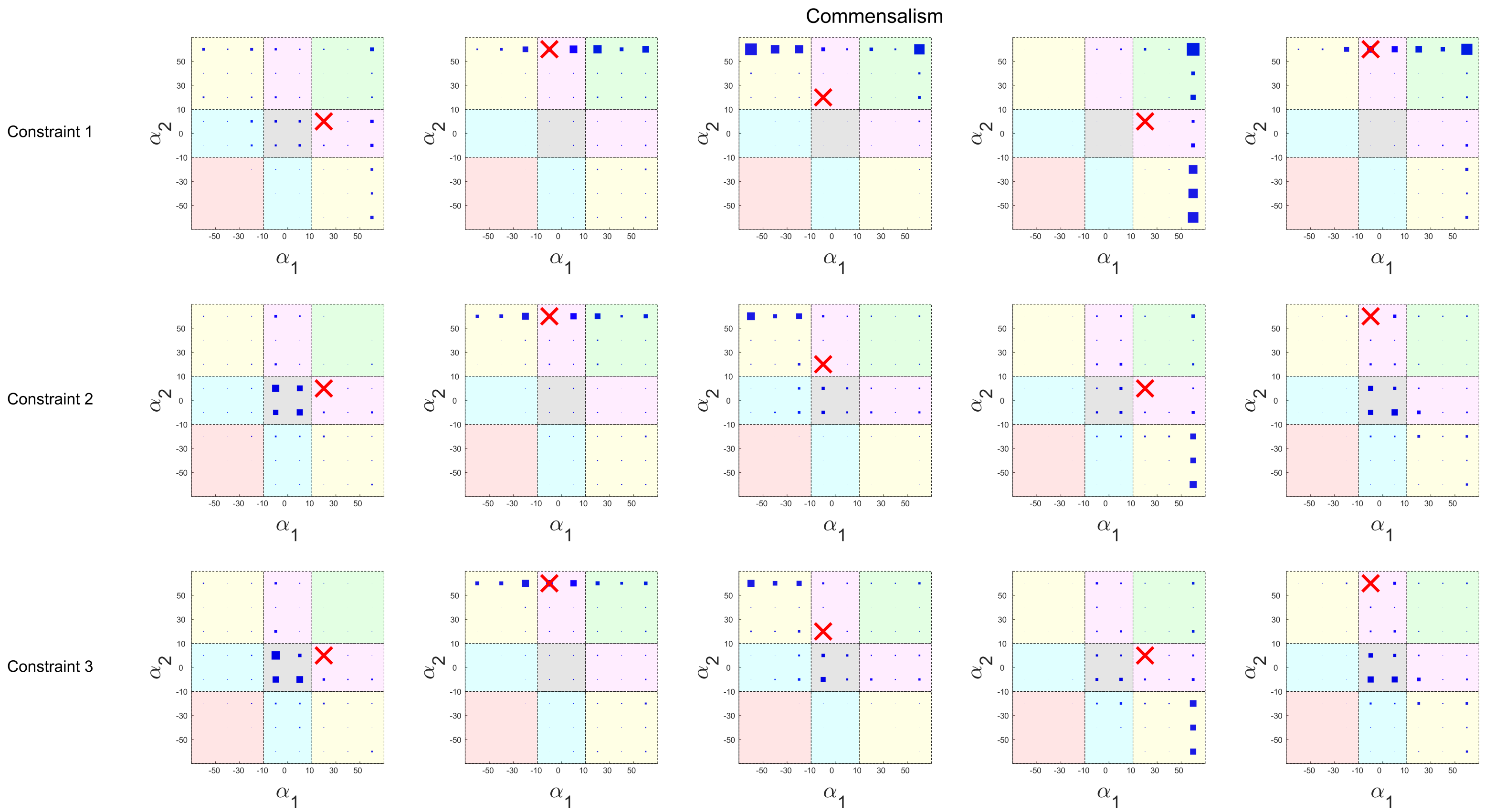

Figure S2: **Heat map of $\alpha_1$ and $\alpha_2$ values for competition starting communities.** The figure depicts the density of $\alpha_1$ and $\alpha_2$ in each bin for all the evolved networks from a starting community that exhibits competitive behaviour. Each sub-graph denotes the evolved communities from an example starting community. The different rows represent the constraints. The graph is coloured into regions based on the cut-off of 10% into various interaction types as green-mutualism, pink-commensalism, yellow-parasitism, blue-amensalism, grey-neutralism, orange-competition. The red cross denotes wild-type starting community.

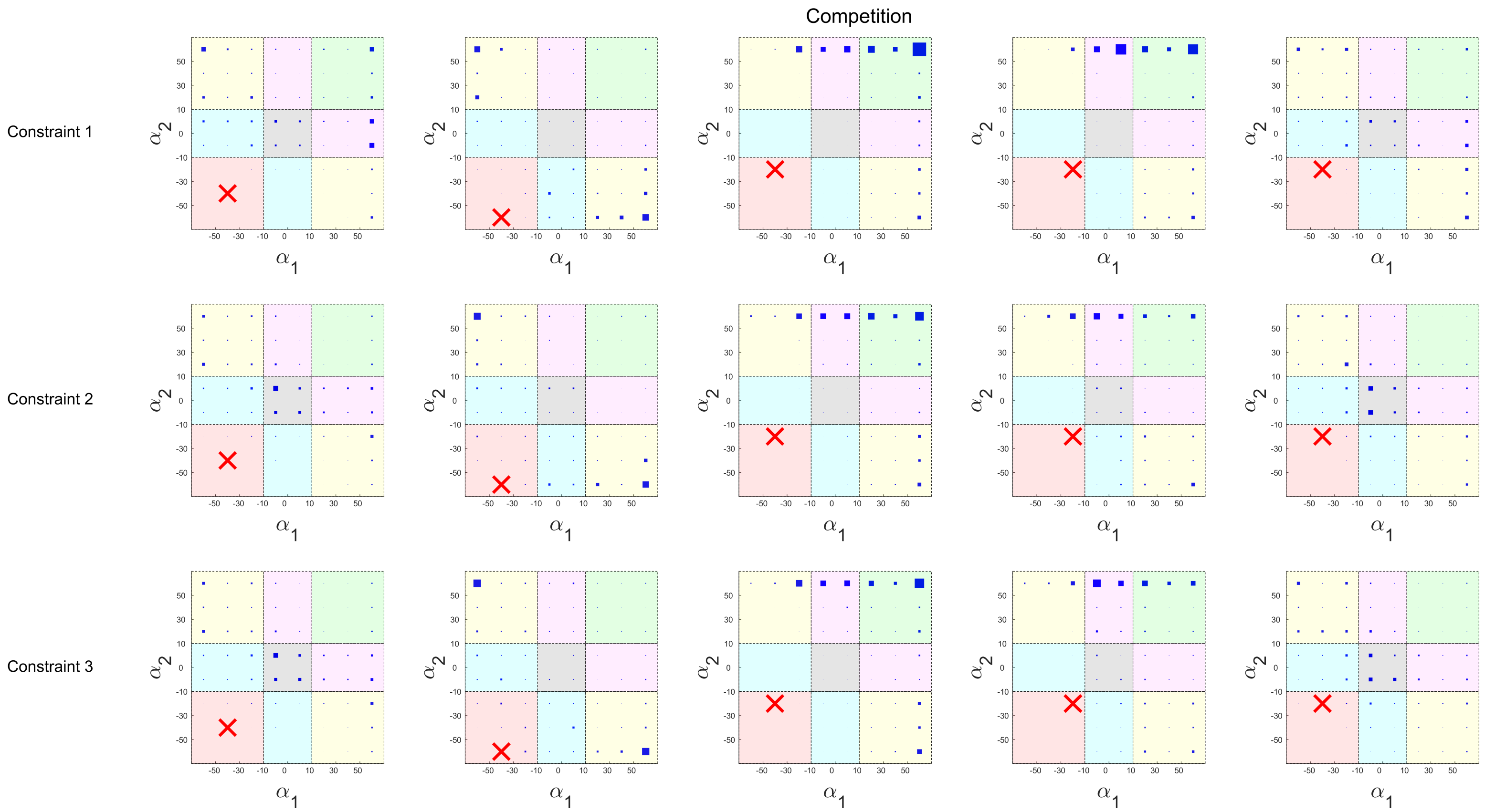

Figure S3: **Heat map of $\alpha_1$ and $\alpha_2$ values for mutualistic starting communities.** The figure depicts the density of $\alpha_1$ and $\alpha_2$ in each bin for all the evolved networks from a starting community that exhibits mutualistic behaviour. Each sub-graph denotes the evolved communities from an example starting community. The different rows represent the constraints. The graph is coloured into regions based on the cut-off of 10% into various interaction types as green-mutualism, pink-commensalism, yellow-parasitism, blue-amensalism, grey-neutralism, orange-competition. The red cross denotes wild-type starting community.

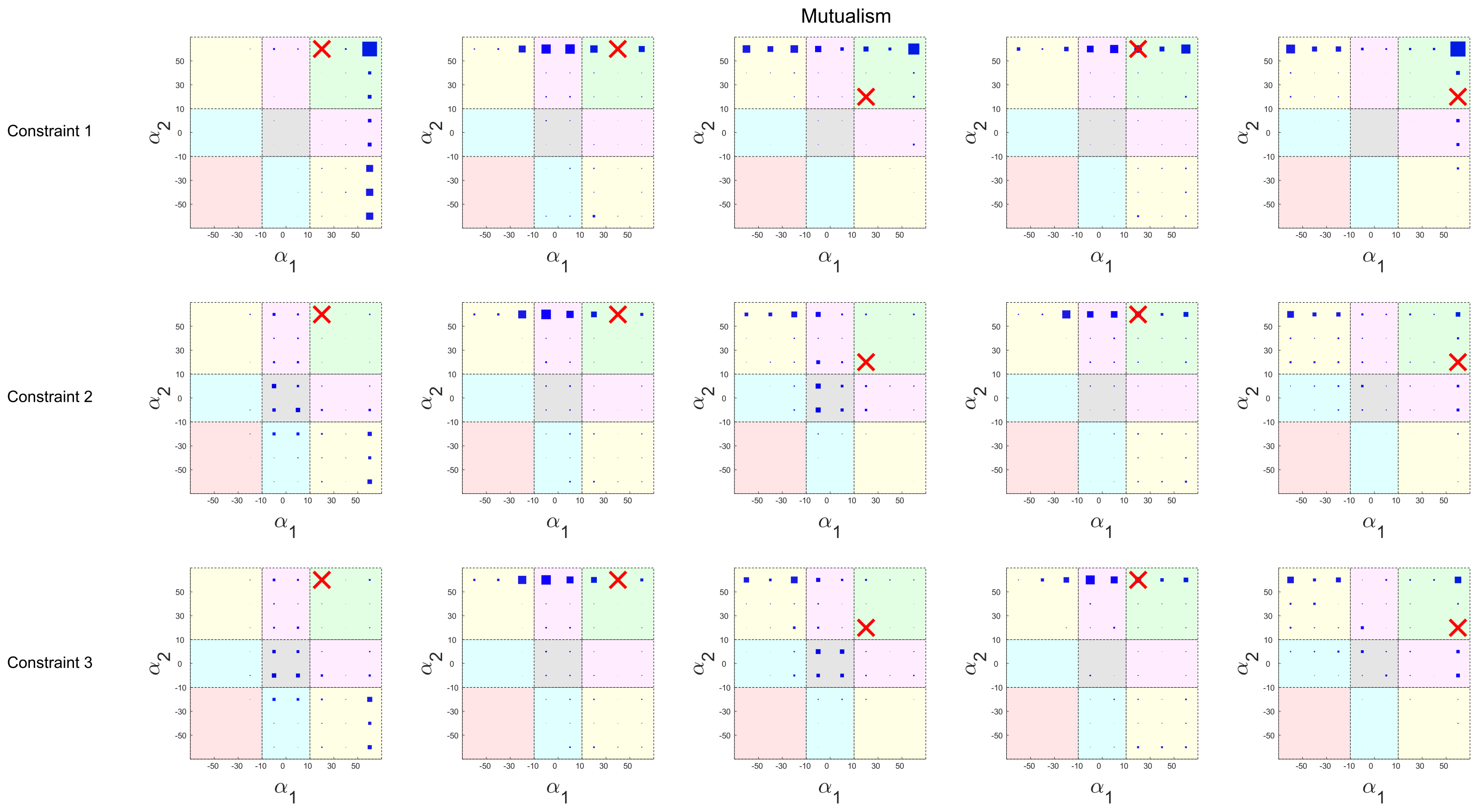

Figure S4: **Heat map of $\alpha_1$ and $\alpha_2$ values for neutralistic starting communities.** The figure depicts the density of $\alpha_1$ and $\alpha_2$ in each bin for all the evolved networks from a starting community that exhibits neutralistic behaviour. Each sub-graph denotes the evolved communities from an example starting community. The different rows represent the constraints. The graph is coloured into regions based on the cut-off of 10% into various interaction types as green-mutualism, pink-commensalism, yellow-parasitism, blue-amensalism, grey-neutralism, orange-competition. The red cross denotes wild-type starting community.

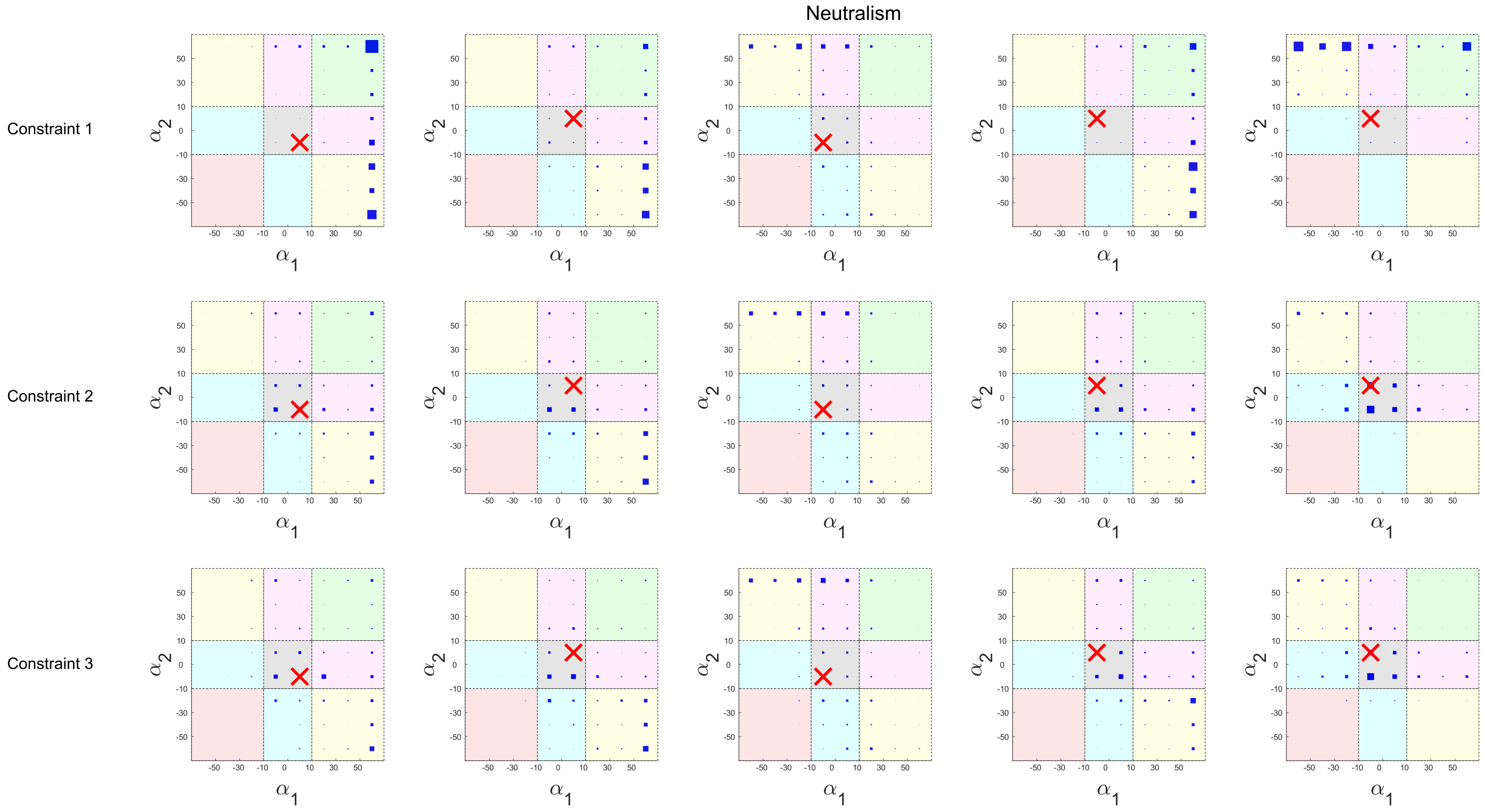

Figure S5: **Heat map of $\alpha_1$ and $\alpha_2$ values for parasitic starting communities.** The figure depicts the density of $\alpha_1$ and $\alpha_2$ in each bin for all the evolved networks from a starting community that exhibits parasitic behaviour. Each sub-graph denotes the evolved communities from an example starting community. The different rows represent the constraints. The graph is coloured into regions based on the cut-off of 10% into various interaction types as green-mutualism, pink-commensalism, yellow-parasitism, blue-amensalism, grey-neutralism, orange-competition. The red cross denotes wild-type starting community.

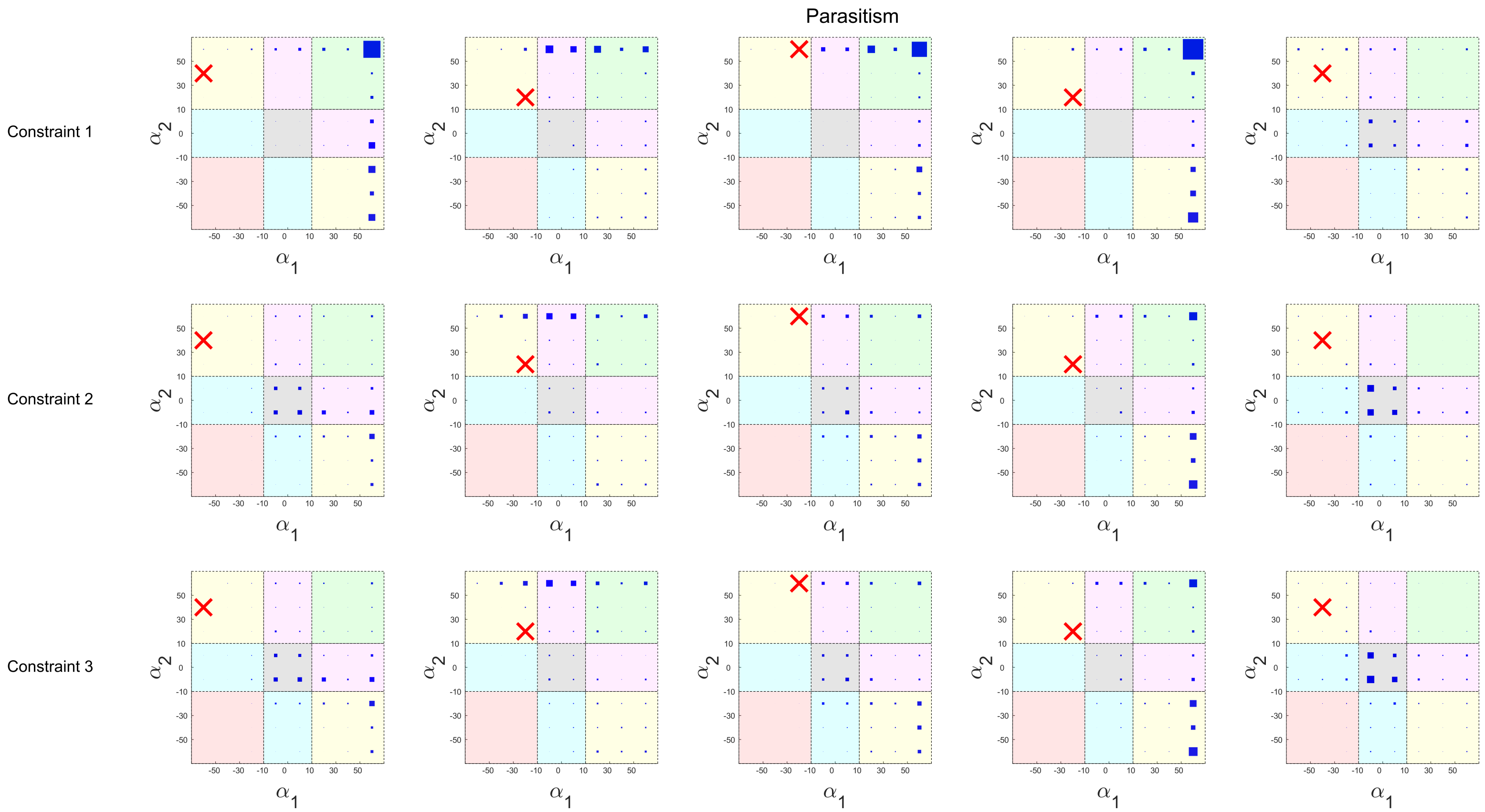

Table S2: **Metabolites that are maximally cross-fed in evolved networks**. The table depicts the metabolites that are maximally cross-fed across the 1000 networks evolved from each example. They have been identified for each wild-type interaction type and documented below.

| Interaction | Example 1 | Example 2 | Example 3 | Example 4 | Example 5 |
|---|---|---|---|---|---|
| **Amensalism** | succ | lys_L | leu_L | glu_L | lys_L |
| **Commensalism** | lys_L | succ | asp_L | lys_L | lys_L |
| **Competition** | leu_L | lys_L | gcald | succ | lys_L |
| **Mutualism** | glu_L | lys_L | asp_L | succ | lys_L |
| **Neutralism** | lys_L | succ | succ | lys_L | glu_L |
| **Parasitism** | lys_L | succ | orn | asp_L | lys_L |
